# Shade-induced WRKY transcription factors restrict root growth during the shade avoidance response

**DOI:** 10.1101/2021.06.01.446656

**Authors:** Daniele Rosado, Amanda Ackermann, Olya Spassibojko, Magdalena Rossi, Ullas V. Pedmale

## Abstract

Shade-intolerant plants rapidly elongate their stems, branches, and leaf stalks to compete with their neighboring vegetation to maximize sunlight capture for photosynthesis. This rapid growth adaptation, known as the shade avoidance response (SAR), comes at a cost; reduced biomass, crop yield, and root growth. Significant progress has been made on the mechanistic understanding of hypocotyl elongation during SAR; however, the molecular account of how root growth is repressed is not well understood. Here, we explore the mechanisms by which low red:far-red induced SAR restrict the primary and lateral root (LR) growth. By analyzing whole-genome transcriptome, we identified a core set of shade-induced genes in the roots of *Arabidopsis* and tomato seedlings grown in the shade. Abiotic and biotic stressors also induce many of these shade-induced genes and are predominantly regulated by the WRKY transcription factors. Correspondingly, a majority of the *WRKY*s were also among the shade-induced genes. Functional analysis using transgenics of these shade-induced *WRKYs* revealed their role is essentially to restrict primary root and LR growth in the shade, and captivatingly, they did not affect hypocotyl elongation. Similarly, we also show that ethylene hormone signaling is necessary to limit root growth in the shade. Our study proposes that during SAR, shade-induced WRKY26, 45, and 75, and ethylene reprogram gene expression in the root to restrict its growth and development. The reduced growth of root organs helps the plant divert its critical resources to the elongating organs in the shoot to ensure competitiveness under limiting photosynthetic radiation.

**One sentence summary:** Shade represses root growth by inducing WRKY transcription factors.

## Introduction

Plants are exposed to various environmental challenges throughout their life cycles, such as suboptimal access to sunlight, low water, nutrient availability, extreme temperatures, presence of competitors, herbivores, and pathogens (Casal, 2012). Plants exhibit incredible plasticity to withstand these adverse conditions and respond by locally adapting growth rhythms, metabolism, and reproduction to best adapt to their environment (Kohnen et al., 2016). An excellent example of adaptive phenotypic plasticity is the shade-avoidance response (SAR). In shade-intolerant plants, SAR is triggered when they are in close proximity to other plant competitors or under a canopy by activating a series of morphological changes to maximize sunlight capture and ensure reproductive fitness (Smith, 1982). The characteristic phenotypes of SAR include rapid stem and petiole elongation, leaf hyponasty, accelerated reproduction, apical dominance, and reduced root growth and development (Salisbury et al., 2007; Casal, 2012) (Fig. 1A). The molecular mechanisms controlling gene expression changes leading to the phenotypic alterations in the shoot organs during SAR are well understood in the model plant *Arabidopsis thaliana* (Casal, 2012; Li et al., 2012; Galvão and Fankhauser, 2015; Pedmale et al., 2016). But the impact of shade on the growth of underground root systems and the molecular account leading to this phenomenon are poorly understood.

**Figure 1.**
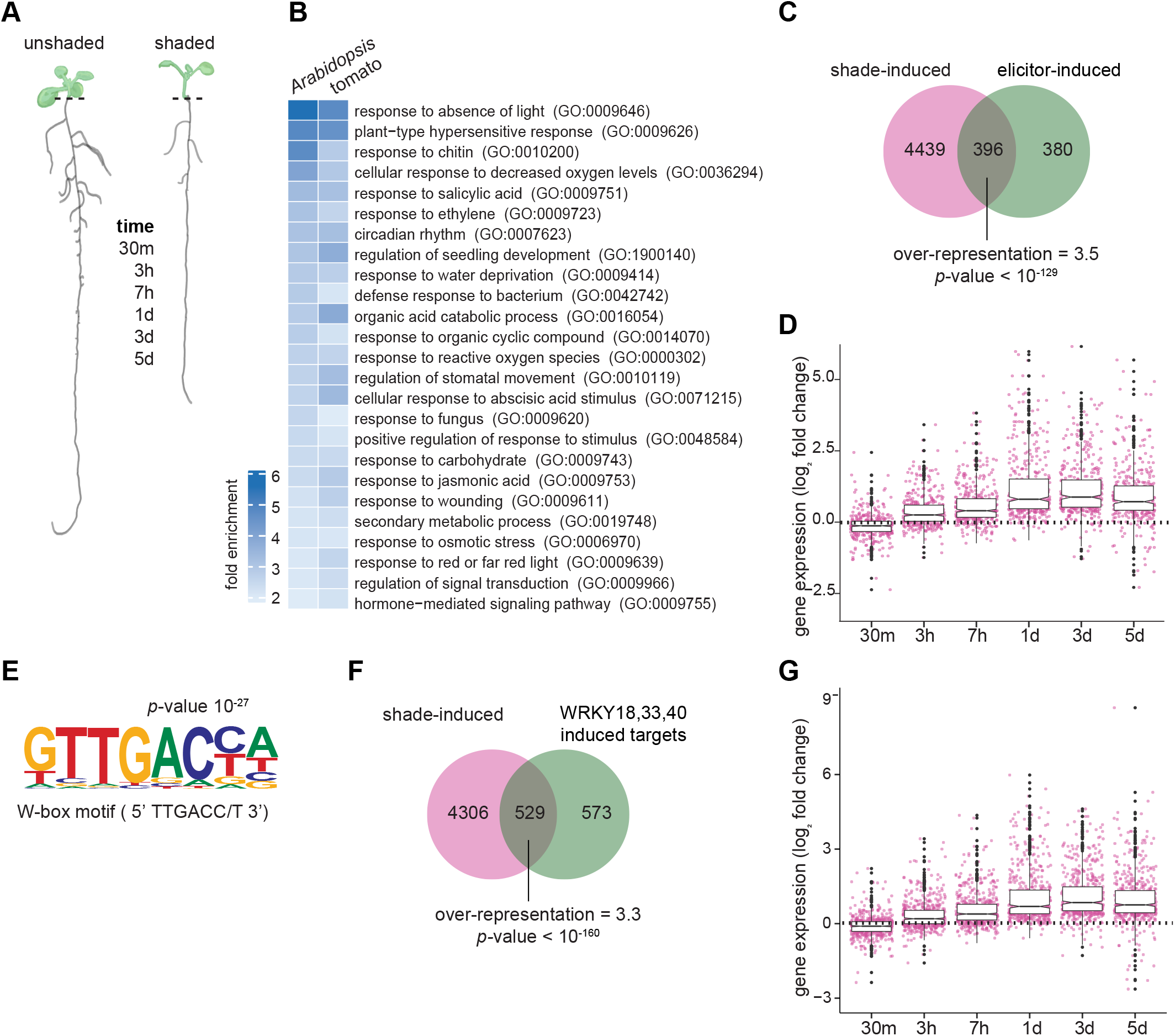
Shade-induced genes in the roots resemble biotic and abiotic stress-induced transcriptome. A) Phenotypic representation of a 9-day-old *Arabidopsis* seedling grown under constant white light (unshaded) and shade (low R:FR); and the time-points used for RNA-seq analysis. Dashed lines indicate the region where the root tissue was excised from the shoot for RNA-seq analysis. B) Gene Ontology (GO) terms of biological processes, commonly enriched among the genes upregulated in *Arabidopsis* and tomato roots during the first 24h (30m, 3h, 7h, 1d; and 3h 6h, 12h, 24h, respectively) under shade. C) *Arabidopsis* genes induced by the shade in the roots along the time-course (up to 5d) and by various PTI-elicitors from a prior study (Bjornson et al. 2021). Over-representation and *p*-value were calculated based on hypergeometric distribution and Fisher’s exact test. D) Expression profile of common *Arabidopsis* shade-induced and elicitor-induced genes along the time-course. Values represent log_2_ fold change in the shade relative to unshaded control. E) *de novo* enriched cis-motif element found in the promoters of the 396 genes induced by both shade and elicitors as determined in a previous study (Bjornson et al. 2021). F) *Arabidopsis* genes induced by the shade in the roots along the time-course (up to 5d) and by WRKY18, WRKY33 and WRKY40 in response to immune response flg22 elicitor from a prior study (Birkenbihl et al. 2017). Over-representation and *p*-value were calculated based on hypergeometric distribution and Fisher’s exact test. G) Expression profile of common *Arabidopsis* shade-induced and WRKY-induced genes along the time-course. Values represent log_2_ fold change in the shade relative to the unshaded control.

Under a dense canopy, plants sense vegetational shading by detecting either a reduction in the ratio of red to far-red (R:FR) light, blue light, or photosynthetically active radiation (PAR) (Keller et al., 2011; Keuskamp et al., 2011; Hornitschek et al., 2012). Any changes in the red/far-red light (R: FR) and blue light in the environment are largely perceived by the R/FR light-sensing phytochrome B (PHYB) and UV-A/blue light-sensing cryptochrome (CRY) 1 and 2 photoreceptors, respectively. In seedlings, CRY- and PHY-mediated shade perception induces the expression of growth-promoting genes in the hypocotyl, such as those involved in hormone biosynthesis and cell-wall remodeling proteins and enzymes, which are both required for the rapid stem elongation (Kohnen et al., 2016; Pedmale et al., 2016; Wit et al., 2016; Paik et al., 2017).

A handful of studies have linked root growth and development with the SAR, and those have mainly focused on the lateral root (LR) emergence and development (Salisbury et al., 2007; Chen et al., 2016; Gelderen et al., 2018; Gelderen et al., 2021). Salisbury et al. (2007) showed that mainly PHYB induces LR formation via auxin signaling and suggested that the inhibition of LR number under low R:FR might be caused by decreased auxin transport or responsiveness in the roots. Findings from Chen et al. (2016) demonstrated that LR development is induced by shoot illumination regardless of the light conditions in which the roots are cultivated, suggesting that a long-distance signal produced in the shoots causes LR formation. It was suggested that ELONGATED HYPOCOTYL 5 (HY5) transcription factor (TF), which is stabilized in the shoots under shade (Pacín et al., 2016), is transported to the roots, where it induces its own expression and regulates LR formation. Based on this observation, van Gelderen et al. (2018) demonstrated that HY5 locally represses LR development in the shade by controlling auxin-dependent pathways at the LR primordia. In a recently published study, it was reported that the expression of hypocotyl-localized HY5 was insufficient to complement the LR growth defects seen in *hy5* mutant *Arabidopsis* plants (Burko et al., 2020).

SAR imparts an important adaptative function to a plant under suboptimal conditions by allowing plants to compete for light. However, such adaptation comes at a cost. For instance, plants prioritize rapid stem and petiole elongation over immunity defense response to herbivores in the shoot, and thus shaded plants are more susceptible to microbial diseases and herbivory (Ballaré, 2014). This prioritization of growth responses over defense is likely to make use of the limited resources efficiently. The presence of pathogens or herbivores activate pattern-recognition receptors present on the cell surface to activate pattern-trigger immunity (PTI), which leads to the induction of salicylic acid (SA) and jasmonic acid (JA)-mediated pathways as a defense response (Ballaré et al., 2012). It has been demonstrated that defense responses including JA signaling are lowered in *phyB* mutant and WT plants exposed to low R:FR shade (Leone et al., 2014; Ortigosa et al., 2020).

Plant disease resistance or biotic stress and abiotic stress responses are primarily mediated by WRKY TFs (Pandey and Somssich, 2009). They constitute the largest family of plant-specific transcriptional regulators, acting as either repressors or activators (Bakshi and Oelmüller, 2014). Accumulating evidence shows that a large number of *WRKY* genes take center stage to regulate various aspects of plant innate immunity by responding to herbivores, PTI elicitors, regulation of defense-related SA and JA hormones, synthesis of defense-related compounds, and phytoalexins (Chi et al., 2013). Apart from their role in stress responses, WRKYs also have diverse biological functions in many plant processes not limited to nutrient homeostasis, seed and trichome development, embryogenesis, seed dormancy, senescence, etc. (Eulgem et al., 2000; Skibbe et al., 2008; Mao et al., 2011; Birkenbihl et al., 2018; Karkute et al., 2018; Viana et al., 2018; Chen et al., 2019). WRKY proteins are largely defined by the presence of a conserved WRKY DNA-binding domain defined by the WRKYGQK amino acid sequence. Apart from the WRKY domain, these transcription factors contain an atypical zinc-finger domain in their carboxyl-terminal (Rushton et al., 2010; Chen et al., 2019). WRKY TFs primarily bind to the W-box cis-elements in the promoter of their target genes (Ciolkowski et al., 2008; Rushton et al., 2010). Thus, WRKYs are essential regulators in responding to internal and external developmental signals as well as stresses.

To understand the molecular account of how low R:FR shade leads to the inhibition of primary root growth, we analyzed the whole genome transcriptome of the roots of *Arabidopsis* and tomato seedlings grown in the shade. We identified a core set of shade-induced genes in the roots of shaded plants, and most of them were also induced by abiotic and biotic stressors. The majority of the shade-induced genes contain W-box promoter elements and are considered the targets of WRKYs. Many *WRKY* gene family members were also significantly upregulated in the roots of shaded plants. To decipher the contribution of individual WRKYs in controlling root growth during the SAR, we overexpressed in *Arabidopsis* a large number of shade-induced WRKYs. We identified that *WRKY26* and *WRKY45* overexpression led to a constitutive-shaded, short primary root phenotype even in the absence of shade. In contrast, overexpression of *WRKY75* lead to a decrease in the LR number in the shade but did not affect the primary root growth. Interestingly, the overexpression of these WRKYs affected only the roots, and it did not lead to any hypocotyl elongation defects seen during the SAR. Similarly, like WRKYs, our study implicates ethylene hormone to be necessary to limit root growth but was insignificant for hypocotyl growth in the shade. In summary, we found that low R:FR shade induces a large number of WRKY TFs, particularly to restrict root growth and development. We hypothesize that the reduced growth of root organs helps the plant divert its critical resources to the elongating organs in the shoot to ensure competitiveness under limiting photosynthetic radiation.

## RESULTS

### Shade-induced genes in the roots resemble biotic and abiotic stress-induced transcriptome

To determine how vegetational shade affects root growth and development, we had performed a whole-genome transcriptomic analysis using RNA-seq as a time course on the excised roots of *Arabidopsis* seedlings grown in unshaded (white light) and shaded (low R:FR) conditions (Fig. 1A and Supplemental Fig. S1A). 5-day-old WL grown *Arabidopsis* seedlings were transferred to shade or mock-treated, then their roots were harvested after 30 min, 3h, 7h, 1d, 3d, and 5d of treatment duration. Similarly, we performed a comparable experiment in 7d tomato seedlings (*Solanum lycopersicum*), and the root tissue was harvested from them after 3h, 6h, 12h and 24h. Total RNA was isolated from these root tissues and the whole genome transcriptome analysis (RNA-seq) was performed using short-read sequencing. Gene expression matrices and statistically significant (false discovery rate; FDR <0.05) differentially expressed genes (DEG) were determined by comparing the shade and unshaded samples to its own developmental time point. We identified a total of 3,395 DEGs that were upregulated in *Arabidopsis*, and 2,523 in tomato by combining all the time points until 24h (Supplemental Fig. S1B; Supplemental Table S1). Henceforth, we will refer these upregulated DEG as shade-induced genes. Next, we subjected these shade-induced genes to Gene Ontology (GO) analysis to assign them a biological function. Our GO analysis on the shade-induced genes in the roots was largely enriched and overrepresented with GO terms related to stress responses, defense against pathogens, and innate immune responses in both *Arabidopsis* and tomato (Fig. 1B; Supplemental Table S2).

As our GO analysis revealed that the shade-induced genes were also induced during biotic and abiotic stress, and plant’s defense against pathogens, therefore, we compared our dataset with the publicly available published RNA-seq datasets, especially related to immunity and defense responses. In one of the comparisons, we chose a recent study in *Arabidopsis* that identified 776 common genes that are induced when treated with seven separate elicitors (3-OH-FA, flg22, elf18, nlp20, CO8, OGs and Pep1) of pattern triggered immunity (PTI) (Bjornson et al., 2021). More than half (51%) or 396 of the 776 elicitor-induced genes overlapped with our shade-induced genes (Fig. 1C), representing an enrichment of 3.5-fold over the number of genes that would be expected by random chance (*p*-value < 10^-129^). These 396 genes, commonly induced by shade and elicitors of PTI displayed an increased temporal expression in roots of *Arabidopsis* seedlings exposed to shade (Fig. 1D), suggesting that prolonged exposure to shade activates defense-like responses in the roots in absence of pathogens.

### Promoters of the shade-induced genes contain W-box elements

To obtain further insights on the nature of the genes that are universally responding to shade stimuli in the roots, we sought to identify the conserved cis-elements in the promoters of shade-induced genes. We performed *de novo cis*-motif analysis on the promoter sequences (500 bp upstream and 50 bp downstream) of the transcription start site of the shade-induced genes in *Arabidopsis* (Supplemental Fig. S1B, Supplemental Table S1) as well as those overlapping with the PTI elicitor-induced genes (Fig. 1C). We identified W-box motif [TTGACC/T] as one of the top enriched *cis*-element among the promoters of the shade-induced genes (*p*-value 10^-17^; Supplemental Fig. S1C) as well among the shade and PTI elicitor-induced genes (*p*-value 10^-27^; Fig. 1E). Approximately 33% of the promoters of the shade-induced genes in *Arabidopsis* roots contained W-box motifs. Interestingly, we did not identify the W-box motif in the promoters of the downregulated genes, instead, we identified cis-motifs typically recognized by TCP (Teosinte branched1/Cincinnata/proliferating cell factor) and MYB transcription factors (Supplemental Fig. S1D). Therefore, considering that WRKYs are central to both biotic and abiotic stresses, we hypothesized that they are likely responsible for the induction of stress and defense-related gene expression program that we observed in the roots of shaded plants (Fig. 1B). Consistently, 48% of the genes known to be directly regulated by WRKY18, WRKY33, and WRKY40 transcription factors via binding to their promoters are also induced by shade (Fig. 1F-G) (Birkenbihl et al., 2017). Among them, well characterized defense marker genes, *CYP71A12, MYB51*, and *PIP1* (Lakshmanan et al., 2012; Hou et al., 2014; Birkenbihl et al., 2017), were found to be significantly induced in the roots of *Arabidopsis* under shade (Supplemental Fig. S1E). Combined, these results suggest that the shade induces genes in the roots that are also upregulated when a plant encounters abiotic and biotic stress, and a large proportion of these genes contain W-box promoter elements, which are binding sites for WRKY TFs.

### Large number of WRKY transcription factors are induced in response to shade

The RNA-seq and GO analysis on the shade-induced genes, and the discovery of W-box promoter elements in them, indicated the involvement of WRKY TFs in mediating root responses to shade. To test this hypothesis, first we surveyed the expression of all the *WRKY* genes in *Arabidopsis* and tomato in our transcriptomic data. We found a large number of *WRKY*s were induced along the time-course in response to shade. 33 out of 74 *WRKY*s in *Arabidopsis* were significantly expressed (FDR <0.05) in one or more time points, similarly, 21 out of 83 *WRKY*s were upregulated in tomato (Fig. 2A-B; Supplemental Fig. S2).

**Figure 2.**
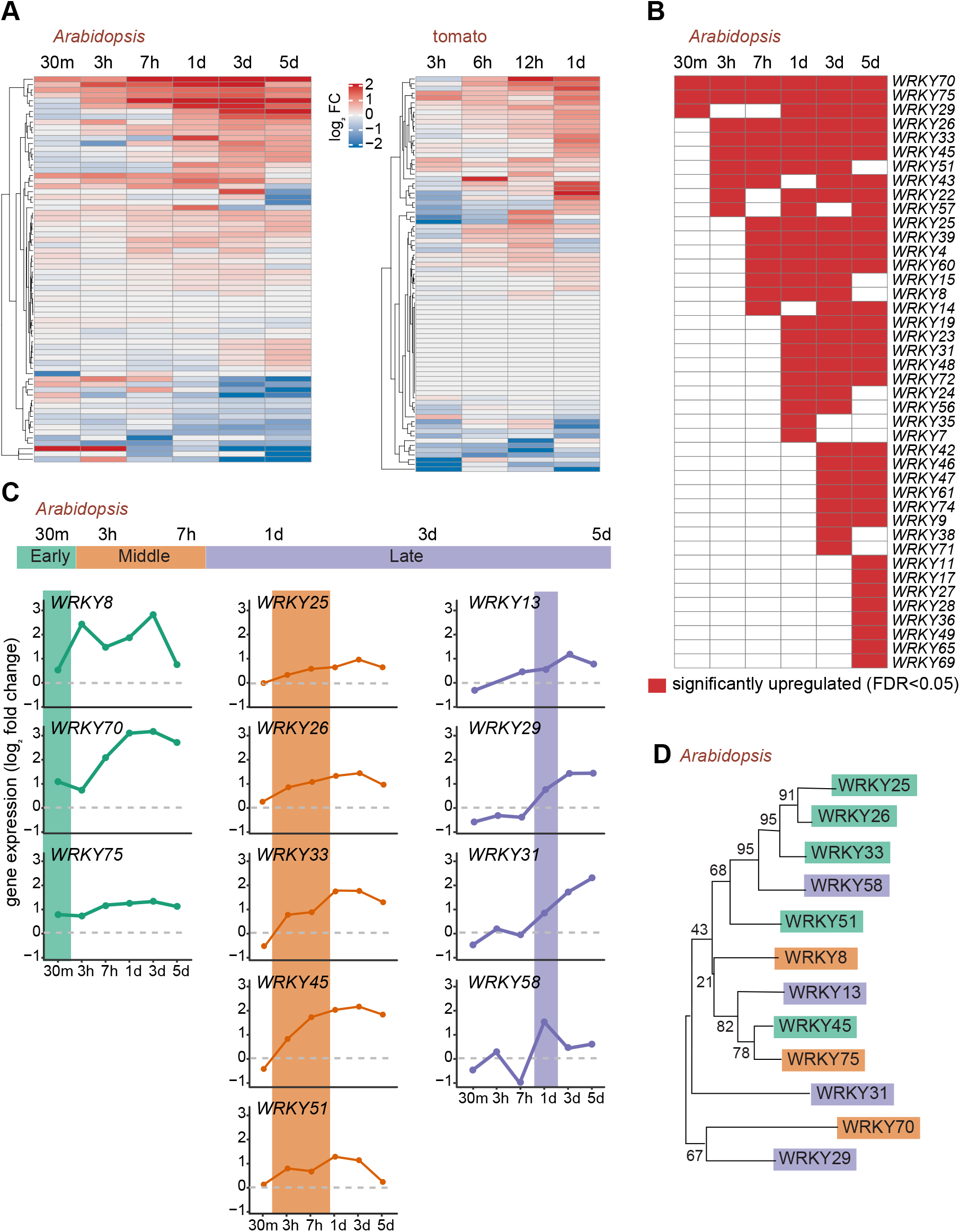
Large number of WRKY transcription factors are induced in response to shade. A) Global transcriptional profile of *WRKY* gene family members in *Arabidopsis* and tomato roots in the shade. Values represent log_2_ fold expression of *WRKY* genes in the shade compared to its unshaded developmental control. B) *WRKY* genes that are significantly up-regulated in the *Arabidopsis* roots relative to its unshaded control (FDR ≤0.05). C) Transcriptional profile of the selected *WRKYs* consistently induced by shade, *i.e*., those up-regulated above the threshold of 0.5 log_2_ fold relative to the unshaded control. Expression groups are color coded and characterized by the timing of their significant induction in the shade during the time course. D) Dendrogram of selected *Arabidopsis* WRKY proteins, classified as early, middle, and late as in panel B. The phylogenetic tree was inferred by Maximum Likelihood and JTT model. Branch lengths are scaled according to number of substitutions per site.

Next, to perform in-depth functional analysis of the shade-induced *WRKY*s in *Arabidopsis*, we sought to narrow down the candidates as large number of them were induced (Fig. 2B). In order to do this, we selected *WRKYs* with a minimum threshold of 0.5 log_2_ fold-change induction relative to the unshaded control along the time-course. Using this parameter, we identified 12 *WRKYs* that were consistently up-regulated in the shade (Fig. 2C). We further classified these 12 genes in to three groups, namely, “early”, “middle”, and “late”, based on the time they were upregulated post shade treatment. *WRKY8, WRKY70*, and *WRKY75* were upregulated within the first 30 min of the shade treatment, there we classified them as immediate early-induced genes. Next, *WRKY25, WRKY26, WRKY33, WRKY45*, and *WRKY51* were classified as intermediate middle, as their expression was seen between 3-7 hours of shading. Lastly, *WRKY13, WRKY29, WRKY31*, and *WRKY58* were classified as late-induced genes as they were expressed only after 24h of shading (Fig. 2C).

Due to the redundancy within the *WRKYs*, we investigated whether the early, middle, and late shade-induced *WRKY* genes were closely related phylogenetically. For this, we constructed a Maximum Likelihood phylogenetic tree based on the amino acid sequences of the 12 WRKY candidates induced by shade. Surprisingly, the pattern of *WRKY* expression did not reflect the phylogenetic relationship between them. For instance, WRKY75 and WRKY45 that are closely related to each other (Fig. 2D), were induced in the middle and at earlier time points (Fig. 2C). An exception to this was the branch comprising WRKY25, WRKY26 and WRKY33, which are phylogenetically close and were expressed in the middle of the time point (Fig. 2C). Also, previous studies have indicated that these WRKYs (25, 26, and 33) act redundantly in *Arabidopsis’* response to high temperature, gibberellin (GA), and abscisic acid (Li et al., 2011; Zhang et al., 2015). Collectively, our results suggest that a large number of WRKY TFs respond to shade stimuli and are specifically induced in both *Arabidopsis* and tomato roots.

### Shade-induced WRKY proteins accumulate in the roots and largely absent in the shoot

As with the known large gene families, various members of the *WRKY* gene family are paralogous, and are documented for functional redundancy due to gene duplications, which complicates genetic analysis to determine the role of individual WRKY TF (Eulgem et al., 2000; Zhang et al., 2015). In this scenario, we decided that the best strategy for studying the role of the shade-induced WRKYs (Fig. 2D) in root growth during shade avoidance is by overexpressing them. It is well documented that overexpression of genes can be used to assess the impact of genetic alterations and gene activity in generating phenotypes (Chua et al., 2006; Prelich, 2012). Therefore, we generated transgenic *Arabidopsis* lines overexpressing the selected shade-induced 12 WRKYs as a mCitrine fluorescent protein fusion (*WRKYox*) under the control of the constitutive *Arabidopsis UBIQUITIN 10* (*UBQ10*) promoter. We identified multiple independent transgenic lines with a single insertion of the transgene and we selected a minimum of three lines for further analysis, except for *WRKY8ox* and *WRKY33ox*, as we could not recover stable transgenic lines for them.

First, we performed immunoblot analysis to ensure that the stable transgenic lines for rest of the ten shade-induced WRKYs were expressing full-length mCitrine fusion proteins, and not partial fusions or free mCitrine alone. Using total protein lysates obtained from the whole 5-day old transgenic seedling grown in unshaded or exposed to the shade for 3-24 hours, we performed immunoblot analysis using an anti-GFP antibody. Immunoblot analysis could detect the presence of full-length mCitrine fusion with WRKYs 25, 26, 31, 45, 51, and 75 among the independent transgenic lines (Supplemental Fig. S3). Differential protein accumulation was not observed in the shade and unshaded growth conditions in these transgenes. However, the specific protein for WRKYs 13, 29, 58, and 70 and some indicated independent lines could not be detected in the immunoblot. This could be due to one or more reasons; transgenic protein expression was below the detection limit of the antibody, low expression of the transgene, dilution of the specific signal due to the use of the whole seedling lysates, or instability of the protein. Nevertheless, we performed confocal microscopy on three independent lines for each of the ten WRKYox-Cit lines that were exposed to a minimum of 24 h of shade. For all of the ten WRKYs, we observed them to be present in the nucleus of the root cells in the maturation zone in shade (Fig. 3A), consistent with their role as a largely nuclear-localized TF (Eulgem et al., 2000). As we had used a constitutive *UBQ10* promoter to express these WRKYs, however, still their protein expression and distribution varied considerably within the cell types of the roots (Fig. 3A; upper panel). In the maturation zone, expression of WRKY29 and WRKY58 proteins were restricted to the outer cell-types, whereas the remaining WRKY proteins were largely observed in all the major cell-types. In the elongation and meristematic zones (Fig. 3A; lower panel), WRKY25ox-Cit, WRKY45ox-Cit, and WRKY51ox-Cit were detected in both the epidermis and cortex, whereas, WRKY13ox-Cit was not detectable. WRKY26ox-Cit was detected only in the columella cells and WRKY58-Cit was observed in the LR cap and the epidermis. In the shoots (Fig. 3B), we detected fluorescence signal for WRKY26-Cit and WRKY45-Cit at very low levels, whereas, WRKY31-Cit signal was observed only in the trichomes. Remarkably, we did not detect any signals for rest of the WRKYox-Cit proteins in the shoot.

**Figure 3.**
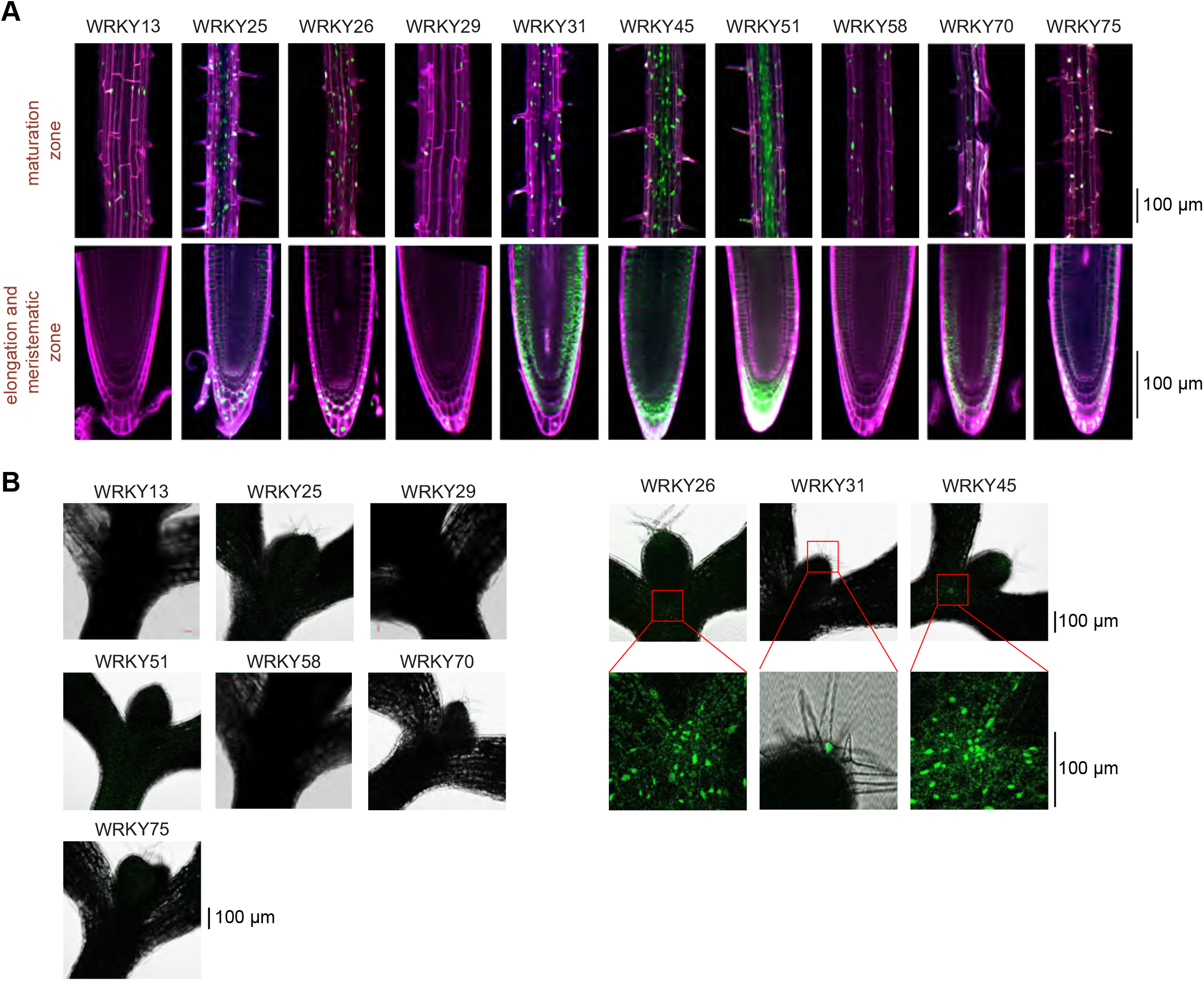
Shade-induced WRKY proteins accumulate in the roots and are largely absent in the shoot. A,B) Confocal microscopy images of the indicated 5-day-old *Arabidopsis* shaded *WRKYox* (fused with mCitrine protein) transgenic seedlings. Prior to the shade treatment, the seedlings were grown in unshaded light for 4 days. A) *Arabidopsis* roots, and B) *Arabidopsis* shoots largely encompassing the hypocotyl, petiole, and the first true leaves. Magenta color indicates the propidium iodide (PI) counterstaining of the cell wall and the green signal indicates mCitrine signal.

Overall, our microscopic analysis revealed the expression of the ten WRKY-Cit proteins, primarily in the roots, but not in the shoots of the transgenic plants in shade, further reinforces the importance of their upregulation in the roots of shaded plants. Also, the expression of these WRKYox-Cit proteins varied, with discrete patterns in different cell-types and developmental zones of the root. This discrete and varied pattern of protein expression which was limited to the roots, likely contributed to the lack of detection of some of the WRKYox-Cit proteins in the are regulated at the protein or gene expression level in a tissue and cell-type specific manner.

### Shade-induced WRKYs affect primary root and lateral root growth in shaded and unshaded conditions

Since many *WRKY*s are upregulated in the roots of *Arabidopsis* and tomato seedlings in the shade, we sought to assess their functional contribution in regulating root growth. We chose to determine the effect of their overexpression in the low R:FR mediated SAR, specifically on hypocotyl elongation and root growth inhibition. For each of the shade-induced selected WRKY overexpressors, we used three independent *Arabidopsis* transgenic lines (described in Fig. 3) for phenotyping, except for *WRKY51*, for which we recovered only two separate lines (Supplemental Fig. S4A). We analyzed four phenotypic traits under unshaded and shade conditions: length of the primary root, LR number, LR density, and hypocotyl length. We report the average phenotypic values in Fig. 4, by combining the measurements from all the independent transgenic lines employed for each of the WRKYs.

**Figure 4.**
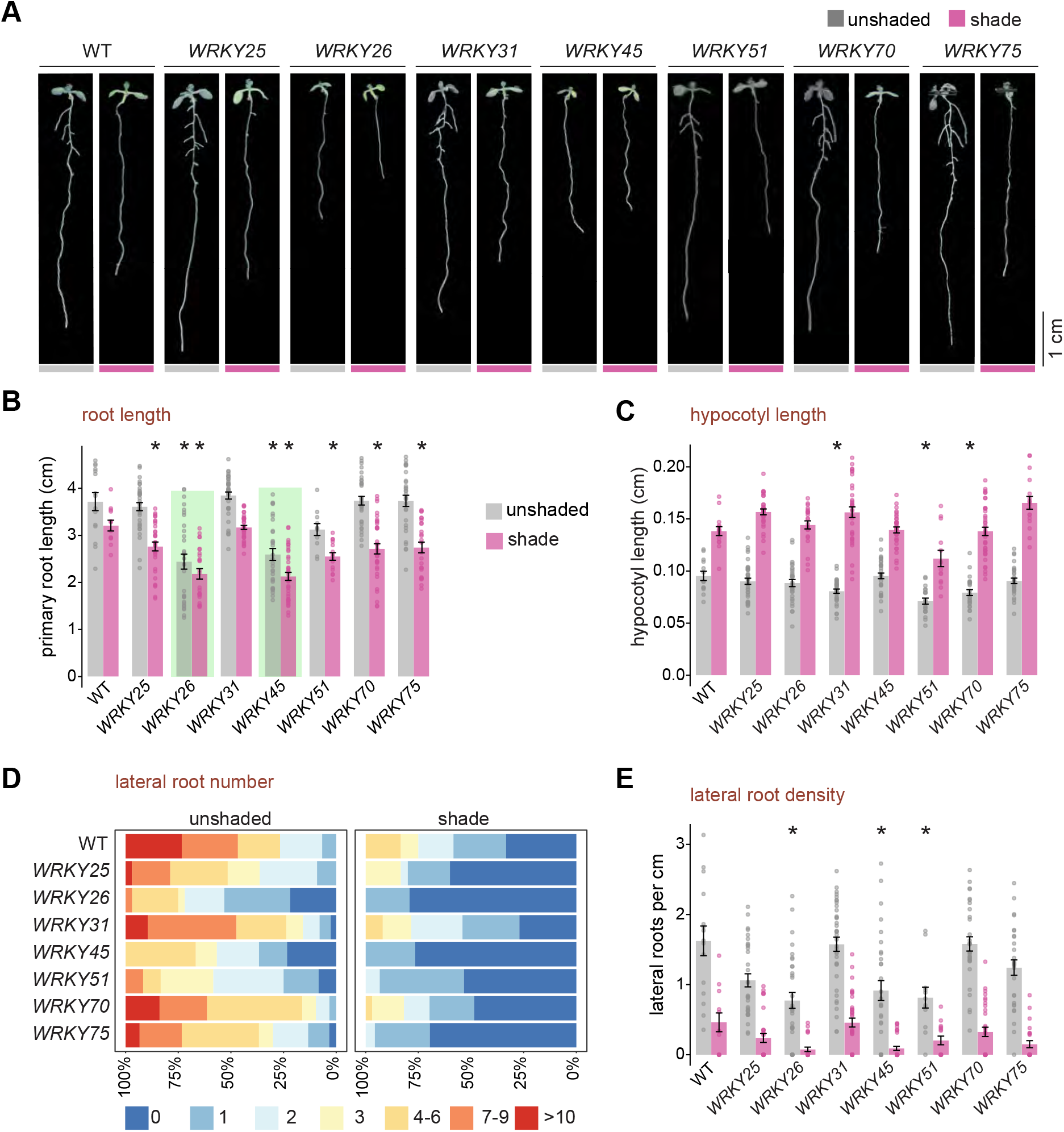
Shade-induced WRKYs primarily affect root growth in the shaded and has no effect on the hypocotyl growth during the SAR. A) Phenotype of representative 9-day-old WT and *WRKY*ox *Arabidopsis* seedlings under unshaded light and shade (low R:FR). B) Primary root length of the indicated genotypes in cm, C) Hypocotyl length of the indicated genotypes in cm, D) Lateral root (LR) density as the number of LR per 1 cm of the primary root. E) Frequency of seedlings with 0-10, or more LRs. The phenotypic measurement represents 4-day-old seedlings grown in unshaded light and then transferred to the shade or mock-treated for 5 days. B-D represent means ± SE of combined independent transgenic lines (*UBQ10::WRKY-mCitrine*) for each candidate gene; dots represent individual data points. Asterisks represent significant difference with WT for each light condition (*p*<0.05) in two-tailed *t*-test with Benjamini-Hochberg correction for multiple testing.

5-day old seedlings grown in unshaded condition were transferred to the shade or mock-treated for 4-days and then their primary root length, LR number, and LR density was measured. Most of the *WRKY* overexpressing seedlings produced primary roots whose length was comparable to the WT (Fig. 4A-B). However, *WRKY26* and *WRKY45* overexpressors displayed a constitutive primary root growth and LR branching inhibition even in the absence of the shade stimuli (Fig. 4A-B). The primary root length of *WRKY26* and *WRKY45* did undergo a modest decrease in size in the shade compared to other *WRKY* overexpressors and the WT (Fig. 4B, Supplemental Fig. S4A).

Surprisingly, all the *WRKY* overexpressing transgenic seedlings did not have any measurable defects in their hypocotyl length and were comparable to the WT (Fig. 4C, Supplemental Fig. S4B) in the shade and non-shading control conditions. *WRKY26ox* and *WRKY45ox*, which displayed hypersensitive shorter roots in non-shading conditions, did not have any obvious hypocotyl growth defects in the shade. However, the expression and accumulation of WRKY26 and WRKY45 protein was detected in the hypocotyl (Fig. 3B). So, the lack of hypocotyl defects in *WRKY26ox* and *WRKY45ox* likely reflects their specialized role in the avoidance response is restricted in regulating limiting root growth.

In all the ten WRKY overexpressing seedlings, we observed reduced LR number in the shaded seedling compared to their unshaded counterparts, similar to WT (Fig. 4D and Supplemental Fig. S4C). In *WRKY26ox* and *WRKY45ox* seedlings, a significantly reduced to nearly absent LRs were noted (Fig. 4A, D), in the shade and non-shading conditions, similar to its constitutive primary root growth inhibition in these conditions (Fig. 4A, B). During the duration of our assay (9 days) in the shade, the LRs were not detected for *WRKY26ox* and *WRKY45ox* seedlings; in contrast, up to 6 could be seen in each WT seedlings (Fig. 4D, Supplemental Fig. S4D). Although WRKY51 and WRKY75 did not influence the inhibition of the primary root growth but, we observed a marked decrease in their number of LR in the shade (Fig. 4D, Supplemental Fig. S4C-D).

Of note, we could only detect a few statistically significant differences in the number of LRs in our transgenic lines (Supplemental Fig. S4C). This is likely due to variation in LR produced by individual seedlings and accounted for by WRKY’s expression levels in the cells. Significant differences were not observed in the LR density, measured as a number of LR per 1 centimeter (cm) of the primary root (Fig. 4E, Supplemental Fig. S3E). This result was not particularly unexpected, considering that most *WRKYox* seedlings did not have much of an impact on the primary root growth and LR number, except for WRKY26, WRKY45, and WRKY75. *WRKY51ox* seedlings also displayed a slight shortening of the primary root and the hypocotyl, but these transgenic lines also showed variable and delayed seed germination, and other pleiotropic defects. Therefore, other confounding factors could be influencing the resulting phenotype of *WRKY51ox* in the shade (Fig. 4A-C).

Overexpression of *WRKY13, WRKY29*, and *WRKY58* showed a slight increase in their root length, LR production, and LR density compared to the WT in the shade and non-shading conditions (Supplemental Fig. S5A-C). However, these observations were reinforced only in the *WRKY13ox* independent lines, but not on *WRKY29ox* and *WRKY58ox* individual seedlings (Supplemental Fig. S5D-F). Considering that these three *WRKY*s are shade-induced in the “late” stages of our time-course analysis (Fig. 2C), it is plausible that their role could be in opposing shade-mediated repression of root growth.

Here, overexpression of several shade-induced WRKYs produced no observable mutant phenotypes with the primary root and LR growth. The lack of phenotypes could be due to the absence of required activating factor, which had to be overexpressed along with the WRKYs. Several WRKYs are also known to be regulated by Ca^2+^ and bind to 14-3-3 proteins (Rushton et al., 2010). Another aspect could be the feedback loops, which could interfere with WRKYs even though they were overexpressed. Several reports point that WRKYs are capable binding to their own promoters or of other *WRKY* genes in response to stress (Skibbe et al., 2008; Rushton et al., 2010; Li et al., 2011; Li et al., 2015; Birkenbihl et al., 2018). Expression and post-translational modification could further limit the function of the shade-induced WRKYs when overexpressed. This is evident from our microscopic analysis (Fig. 3), as several of them were not detected in the shoot and also restricted to certain cell-types in the root. Epitope tags are routinely fused with TFs, and they rarely interfere with the function of the TFs as determined by various *in vitro* and *in vivo* experiments such as ChIP-seq (chromatin immunoprecipitation-sequencing), etc. However, we cannot rule out completely whether the fusion of the WRKYs with mCitrine protein could be interfering with their activity.

Together, our results here indicate the significance of WRKYs, especially the upregulation of *WRKY26* and *WRKY45* in the roots of shaded plants to limit their growth and not that of the hypocotyl. Overexpression of *WRKY26* and *WRKY45* resulted in reduced primary root growth and LR number under the control unshaded condition, as to resemble the phenotype of WT roots in the shade. *WRKY75ox* had no significant role in repressing the primary root growth, but had an affect by repressing LR emergence in the shade and not in unshaded growth.

### Ethylene is required for root growth inhibition in the shade

Apart from mediating responses to biotic and abiotic stress responses, WRKY TFs integrate ethylene hormone responses along with environmental and developmental signals (Koyama, 2014). Furthermore, ethylene and components of the ethylene signaling pathway are required for efficient resistance towards certain plant pathogens. For example, ethylene-insensitive *Arabidopsis* mutant *ein2* was more susceptible than WT plants to infection by *Botrytis cinerea* fungal pathogen (Thomma et al., 1999). Involvement of ethylene signaling and multiple WRKYs in response to senescence and high temperature have been documented (Li et al., 2011; Koyama et al., 2013; Koyama, 2014). Ethylene also regulates root growth by mostly restricting cell elongation (Růžička et al., 2007). Previous studies have determined the importance of ethylene in petiole elongation but not for hypocotyl elongation during SAR (Pierik et al., 2009; Das et al., 2016). Therefore, in light of this prior knowledge, we tested the effect of ethylene and the components of ethylene signaling on root growth during shade avoidance.

We grew WT seedlings in unshaded and shaded conditions on a growth media supplemented with different concentrations of 1-aminocyclopropane-1-carboxylic acid (ACC), a routinely used biosynthetic precursor of ethylene (Růžička et al., 2007). In the shade, a lower dose of 0.2 μM ACC had a stimulatory effect on the hypocotyl elongation. In contrast, higher doses of 2 and 10 μM had a moderate impact on the hypocotyl elongation (Fig. 5A), confirming previous results (Das et al., 2016). But ACC treatment profoundly affected the primary root growth in the seedling grown in both shade and unshaded conditions (Fig. 5B). At 0.2 μM ACC, the primary root growth of unshaded seedlings was indistinguishable from that of shaded seedlings, having a shorter root length which was comparable to untreated roots in the shade. Increasing concentrations of ACC led to further attenuation of the root growth in both unshaded and shade conditions with similar root length, especially at 10 μM of ACC. ACC treatment, notably at 10 μM, led to a modest reduction in LR number in unshaded seedlings (Fig. 5C). However, ACC had the opposite effect on shaded seedlings, as we observed increased LR density at 10 μM ACC (Fig. 5D). At 2 and 10 μM ACC, the LR density in shaded and unshaded seedlings was similar (Fig. 5D). Therefore, these results indicate that ethylene is required for root growth inhibition in the shade.

**Figure 5.**
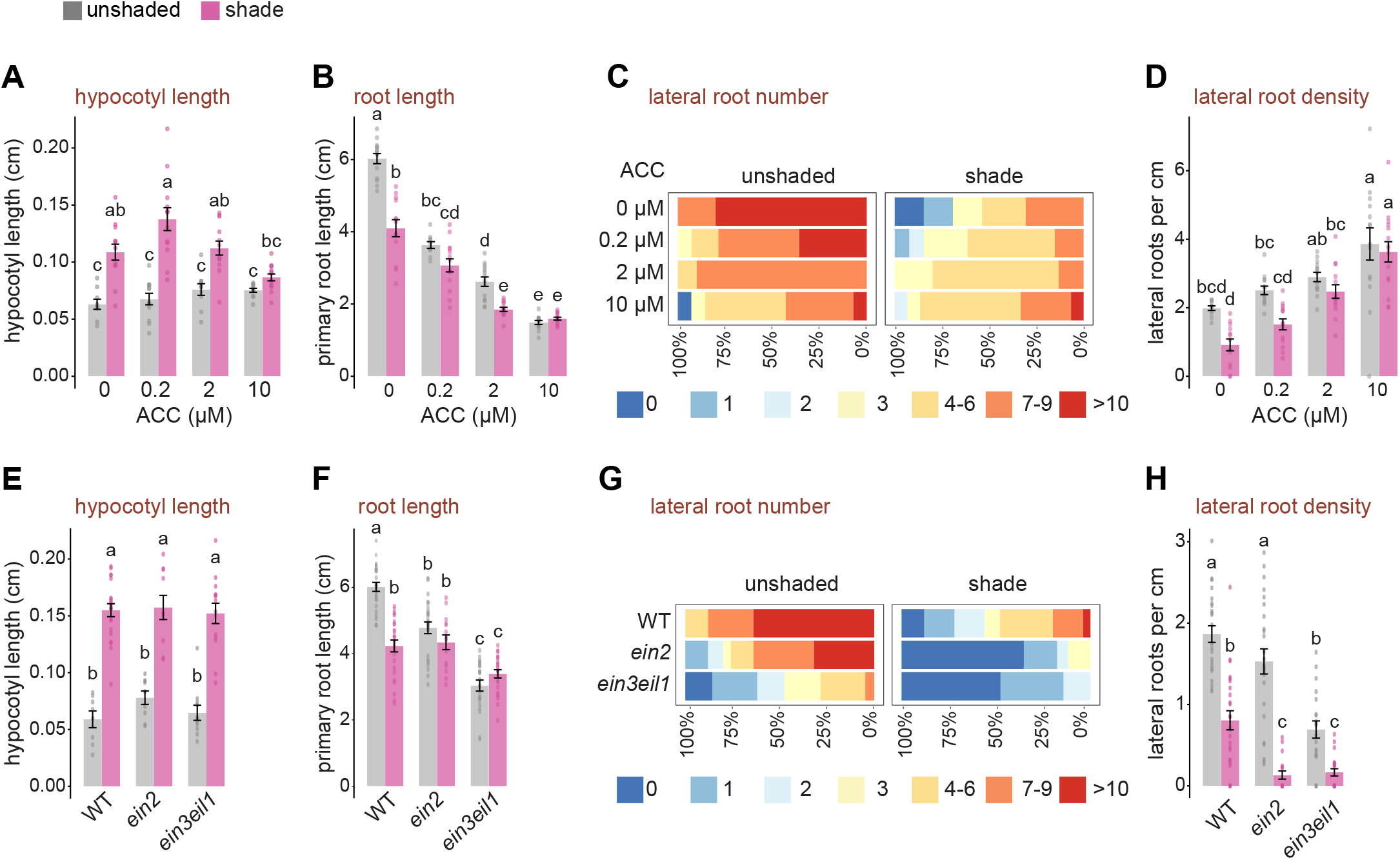
Ethylene hormone signaling is required for the repression of root growth and development in the shade. A-D) Phenotype of unshaded and shaded WT seedlings treated with the indicated concentrations of ACC. E-H) Seedlings of WT and ethylene signaling mutants grown in shaded and unshaded light. The phenotypic measurement represents 4-day-old seedlings grown in unshaded light and then transferred to the shade or mock-treated for 4 days. A, E) Hypocotyl length in cm, B, F), Primary root length in cm, C,G) Frequency of seedlings with 0 – 10, or more lateral roots, D,H) Lateral root density is the number of lateral roots per cm of the primary root. Values represent means ± SE; dots represent individual data points. Different letters denote significant differences in *post-hoc* Tukey test (ANOVA *p*<0.05).

Next, to further explore the importance of ethylene in regulating root growth in the shade, we analyzed mutants defective in ethylene signaling, namely, *ein2* and *ein3eil1* double mutant, respectively. EIN2 (ETHYLENE INSENSITIVE 2) is a crucial signaling transducer, and EIN3 (ETHYLENE INSENSITIVE 3) and EIL1 (EIN3-LIKE 1) are critical downstream transcription factors in the ethylene response (Dolgikh et al., 2019). Hypocotyl growth defect was not observed in *ein2* and *ein3eil1* seedlings in the shade (Fig. 5E), agreeing with a previous study (Das et al., 2016). Interestingly, the primary roots of both *ein2* and *ein3eil1* mutants did not respond to shade-induced growth inhibition, regardless of the shade or unshaded growth conditions (Fig. 5F). The root phenotypes of *ein2* and *ein3eil1* further resembled *WRKY26ox* and *WRKY45ox* seedlings (Fig. 4B-E). Accordingly, *ein2* and *ein3eil1* mutants presented much fewer LRs and LR density compared to the WT, both in the shade and non-shading conditions (Fig. 5G-H). Together, the data presented in Fig. 5 indicates that ethylene and its associated signaling is required to restrict root growth in the shade. Also, the data supports the requirement of ethylene signaling along with WRKYs, analogous with the plant defense responses.

## DISCUSSION

Mechanisms underlying stem and petiole elongation under shade have been widely studied for several decades (Hornitschek et al., 2009; Pierik et al., 2009; Li et al., 2012; Pedmale et al., 2016). However, our understanding of how shade perceived by the above-ground shoots leads to the reduced growth of the belowground primary root and LR has been limited. This study presents evidence that many *WRKY* genes are transcriptionally upregulated in the roots of shaded *Arabidopsis* and tomato plants. We further demonstrate that several WRKYs (26, 45, 75) and ethylene function in restricting root and LR growth but did not affect hypocotyl elongation in the shade.

We discovered genes induced by biotic and abiotic stressors overlapped with a large proportion of the shade-induced genes in the roots of *Arabidopsis* and tomato seedlings grown in the shade. PTI-induced genes also coincided with a significant portion of the shade-induced genes in the roots, which are known to be regulated by WRKY TFs. Importantly we found W-box promoter elements in a large number of shade-induced genes in the roots (Fig. 1E, Supplemental Fig. S1C), suggesting the involvement of WRKYs in the reprogramming of the gene expression to restrict root growth, typically observed during SAR. Furthermore, a significant proportion of *WRKY* gene family members were upregulated in the roots of both shaded *Arabidopsis* and tomato, progressively increasing through the time course in the shade (Fig. 2, Supplemental Fig. S2). To identify the contribution of shade-induced WRKYs in regulating root growth in the shade, we performed functional analysis on a select ten WRKY members by overexpressing them. We chose overexpression as an alternate yet powerful tool to generate mutant phenotypes (Chua et al., 2006; Prelich, 2012) and also to overcome known functional and genetic redundancy among the WRKY gene family members and potential gene-compensation (Rushton et al., 2010; Zhang et al., 2015). Overexpression of *WRKY26* and *WRKY45* led to a retarded root growth and LR emergence, irrespective of shade or unshaded light. *WRKY26ox* and *WRKY45ox* seedlings had a constitutive shade avoiding shorter primary root and reduced LR in unshaded light, mimicking a WT seedling in the shade. Importantly, in *WRKY75ox* seedlings, there was no effect on the primary root length, but a marked reduction in LR number was seen, similar to *WRKY26* and *WRKY45* overexpressors. But, none of the ten shade-induced WRKYs that we characterized affected hypocotyl elongation, indicating that the roles of these WRKYs are primarily limited to regulate root growth during the SAR.

Our results here demonstrate that phenotypic activation of WRKY TFs is feasible as a general approach to identify their functional roles in plant growth and development (Fig. 4). Previous studies have shown enhanced resistance towards pathogens, salt, and drought by overexpressing WRKYs that was under investigation in *Oryza sativa* (rice), *Glycine max* (soybean), and *Arabidopsis* (Rushton et al., 2010). However, apart from this study, only a few prior reports have associated WRKYs with light signaling and adaptation. For instance, *Arabidopsis* WRKY18 and WRKY40 have been shown to co-localize with PHYB and PHYTOCHROME-INTERACTING FACTORS (PIFs) in the nuclear speckles or photobodies, but their role in red/far-red light signaling is unknown (Geilen and Böhmer, 2015). WRKY40 is required for adaptation towards high light stress in *Arabidopsis* (Aken et al., 2013), and WRKY22 is involved in dark-induced senescence (Zhou et al., 2011).

Previously, WRKY26 was identified as a positive regulator of thermotolerance in *Arabidopsis* plants, working synergistically with *WRKY25, WRKY33*, and ethylene signaling (Li et al., 2011). In this literature, overexpression of *WRKY26* led to the reduction of fresh weight in adult *Arabidopsis* plants, mirroring our results, where its overexpression led to shorter roots in the shade and unshaded light (Fig. 4). *Arabidopsis* WRKY45 is a positive regulator of GA-mediated age-induced leaf senescence (Chen et al., 2017), and has been implicated in the activation of *PHOSPHATE TRANSPORTER1;1 (PHT1;1)* in the roots of plants undergoing phosphate deficiency (Wang et al., 2014). Wang et al. (2014) further showed that *WRKY45ox* lines had shorter primary roots in the presence of arsenate. This result is similar to our study, where *WRKY45ox* led to constitutive primary root length shortening in the shade and unshaded light. Furthermore, we report that *WRKY75* overexpression had no effect on the primary root length but caused a reduction in lateral root frequency in seedlings grown only in the shade but not in unshaded conditions (Fig. 4D and Supplemental Fig. S4D). Silencing of *WRKY75* by RNAi resulted in more LR than WT *Arabidopsis* but no differences in the primary root length (Devaiah et al., 2007). Also, WRKY75, like WRKY45, can promote leaf senescence, participating in a positive feedback loop with hydrogen peroxide and SA to accelerate leaf senescence (Guo et al., 2017). WRKY75 is also a positive regulator of GA-mediated control of flowering time (Zhang et al., 2018). Surprisingly, although WRKY45 and WRKY75 have overlapping roles in phosphate acquisition, leaf senescence, and GA-mediated signaling, yet, only *WRKY45ox* seedlings displayed inhibitor of root length, indicating that many WRKYs share many overlying as well as divergent functional roles.

Ethylene is necessary for petiole elongation during the SAR but had a limited effect on hypocotyl elongation (Pierik et al., 2009; Das et al., 2016). Our results (Fig. 5) show that ethylene and its associated signaling are required to restrict root growth in the shade, suggesting an organ-specific role of ethylene signaling in the shade. Ethylene signaling is critical for a plants’ defense responses and molecular links between this hormone and WRKY activity have been proposed (Bakshi and Oelmüller, 2014). Ethylene induces the expression of several *WRKY*s, and WRKY25, 26 and 33, induce the expression of *EIN2*, forming a positive feedback loop (Li et al., 2011; Li et al., 2013). Expression of *EIN3* is induced by ethylene via EIN2, and also by the defense-hormone JA, and by light via PIF4 and PIF5 bHLH transcription factors (Li et al., 2013; Sakuraba et al., 2014). Our data showed that ethylene-treated WT seedlings and *ein2* and *ein3eil1* had similar root phenotypes to *WRKY26ox* and *WRKY45ox* lines in the shade (Fig. 4, Supplemental Fig. S4, Fig. 5). WRKYs can function up, and downstream of the many phytohormone pathways (Antoni et al., 2011). Therefore, it is conceivable that ethylene and the shade-induced WRKYs act in concert to antagonize root growth and development and signaling dependent upon auxin, brassinosteroid, and cytokinin hormones in the shade.

WRKYs are plant-specific TFs and are central components of the plant’s resistance to pathogens and responses to abiotic and biotic stresses (Bakshi and Oelmüller, 2014). *Arabidopsis* and tomato genomes have 74 and 83 WRKY genes, respectively, and a great diversity of WRKYs allows plants to cope with various adverse conditions (Bakshi and Oelmüller, 2014). WRKYs have been well studied in their involvement with biotic stress compared to abiotic responses (Bakshi and Oelmüller, 2014). SAR is also stressful for the plant, as it leads to photosynthetic impairment and reduced carbon acquisition, reducing the overall fitness of the plant (Smith, 1982; Smith, 2000). In addition, low R:FR SAR downgrades plant defense against pathogens and herbivorous insects in the shoots (Moreno et al., 2009; Ballaré et al., 2012; Courbier and Pierik, 2019). Likewise, other studies have shown that many *WRKY* genes are rapidly induced in response to wounding, drought, salinity, osmotic, cold, carbon starvation, and heat stress (Pandey and Somssich, 2009; Bakshi and Oelmüller, 2014; Rinerson et al., 2015; Viana et al., 2018). Aptly, our finding on the speedy upregulation of *WRKY* gene expression in the roots of shaded plants is conceivable. During SAR, stress-like responses likely help the plant to relocate critical resources from the root to the growing shoot organs in order to maintain competitiveness. Future studies will be required to determine the nature of the signal downstream of the photoreceptors that leads to the induction of many *WRKY* genes in the shade. Also, which are the specific gene targets of the shade-induced *WRKY*s required to repress root growth and development in the shade?

### Conclusions

Low R:FR shade represses root growth and development by triggering gene expression changes in tomato and *Arabidopsis*, mainly resembling the transcriptional changes caused by biotic and abiotic stressors. These shade-induced genes in the roots are known to be regulated by WRKY transcription factors, and many *WRKY*s were significantly upregulated in the shade. Here, we report the involvement of crucial WRKY TFs, namely WRKY26, WRKY45, and WRKY75, along with ethylene in the SAR to inhibit primary root and LR growth. These factors had no involvement in regulating hypocotyl elongation.

## MATERIALS AND METHODS

### Plant Material, growth conditions and light treatments

*Arabidopsis thaliana* lines used in this work are in Columbia (Col-0) ecotype background. Seeds of *ein2-5* and double mutant *ein3-1 eil1-1* were obtained from Dr. Hong Qiao (University of Texas at Austin). For phenotyping of seedlings, seeds were surface sterilized with 70% ethanol and 0.1% triton and rinsed several times with sterile water. Seeds were plated on 0.5× Linsmaier and Skoog (LS) Medium (HiMedia Laboratories) pH 5.8 containing 0.8% phyto agar. Plates with the seeds were then stratified at 4°C for 4-5 d before being placing them vertically in the growth chamber with constant white light (unshaded) at 23°C. After 4 d, plates were either kept under white light (unshaded control) or transferred to simulated shade (R:FR = 0.35) for 1d (microscopy and immunoblot analysis) or 4-5d (phenotypic measurements). For phenotypic measurements, plates with 9 d-old seedlings were scanned using a flatbed scanner (Epson V600), primary root length and later root (LR) numbers (counted as emerged LR) were obtained with SmartRoot (Lobet et al., 2011) and hypocotyl length was measured using NIH ImageJ software.

### Cloning and generation of transgenic lines in *Arabidopsis*

*Arabidopsis* root tissue cDNA library was used as a template to amplify and clone the coding sequence of *WRKY13, WRKY25, WRKY26, WRKY29, WRKY31, WRKY45, WRKY51, WRKY70* and *WRKY75*; while *Arabidopsis* genomic DNA was used as a template to amplify *WRKY58*. PCR was performed using KOD polymerase (Toyobo) with the primers listed in Supplemental Table S3 and introduced in to the Gateway donor vectors, either pDONR207 or pDONR221 (Thermo Fisher Scientific) using BP clonase II enzyme (Thermo Fisher Scientific). Multisite Gateway reaction using LR clonase II mix (Thermo Fisher Scientific) was performed to combine the donor constructs with either pB7m34GW or pK7m34GW binary vector (Karimi et al., 2007) along with the *UBQ10* promoter in pDONR P4-P1R donor vector and Citrine in pDONR P3-P1R vector (destination vectors used for each *WRKY* gene are listed in Supplemental Table 2). Destination constructs were introduced in *Agrobacterium tumefaciens* to transform *Arabidopsis* using the floral dip method (Clough and Bent, 1998). T1 transgenic *Arabidopsis* plants were selected on 0.5× LS medium supplemented with either kanamycin or basta according to the vector used for transformation (Supplemental Table S3). Segregation analysis was performed on T2 plants grown on the selective agar media and lines carrying a single copy of the transgene was propagated further and the T3 seeds were used for experiments and other analysis.

### Immunoblot analysis of transgenic lines

Seedlings were grown constant white light for 4d and either kept in unshaded conditions of shade for additional 24h. 20 excised roots or whole-seedlings were harvested from 5-day old plants and immediately frozen in liquid nitrogen. Samples were ground to a fine powder, resuspended in 2× LDS sample buffer (53mM Tris-HCl, 70.5 mM Tris base, 1% LDS, 5% glycerol, 0.255mM EDTA, 0.11mM Serva Blue G250, 0.0875 mM phenol red, pH 8.5) with 50 mM TCEP and heated to 90°C for 10 min, cooled to room temperature and then centrifuged for 5 minutes to obtain the total protein lysate. For electrophoretic separation of proteins, equal amount of total protein was loaded on 10% Bis-Tris polyacrylamide gel and electrophoretically separated using MOPS-SDS buffer (2.5mM MOPS, 2.5mM Tris base, 0.005% SDS, 0.05mM EDTA). Separated proteins were then transferred electrophoretically to a nitrocellulose membrane (GE Lifesciences) using transfer buffer (10% methanol, 1.25mM bicine, 1.25mM Bis-Tris base, 0.05 mM EDTA). After transfer, the membrane was stained with ponceau red, imaged, and blocked with 5% (w/v) non-fat dry milk prepared in Tris-buffered saline with 0.05% Tween-20 (TBST) for 30 min. Next, membranes were incubated for 1h in 1% milk prepared in TBST with anti-GFP antibody (Roche). The blot was washed 3 times with TBST and incubated for 30 min in 1% milk prepared in TBST with anti-mouse HRP conjugate (Bio-Rad). Chemiluminescent detection was performed using SuperSignal West Dura Extended Duration (Thermo) HRP substrate to detect the WRKY-mCitrine fusion protein.

### Confocal microscopy

Transgenic *Arabidopsis* seedlings expressing *UBQ10::WRKY-Citrine* were grown in unshaded light for 4-5 days and transferred to shade for 24 h. Seedlings were stained with 5 μg/mL propidium iodide and then mCitrine and propidium iodide fluorescence were detected in a high-resolution laser scanning confocal microscope (LSM900 with Airyscan2, Zeiss) using 488 and 561 nm lasers along with BP 620/60 emission filter.

### Phylogenetic analysis

Protein sequences of the 12 candidate *Arabidopsis* WRKYs were retrieved from The Arabidopsis Information Resource (TAIR) database (Garcia-Hernandez et al., 2002). Sequences were aligned with Clustal algorithm, and the phylogenetic tree was inferred by using the Maximum Likelihood method and JTT matrix-based model in Mega X software (Kumar et al., 2018).

### Short-read RNA-seq data

The raw short-read sequencing data and expression files are available in the NCBI Gene Expression Omnibus (GEO) database with accession number GSE175963.

### *De novo* identification of cis-motifs

De novo *cis*-motifs in the promoters of differentially expressed genes were identified with HOMER (Heinz et al., 2010) by analyzing the 500 bp upstream and 50 bp downstream of the transcriptional start site.

### Computational analysis

Gene Ontology (GO) term enrichment was performed on Panther Classification System (Mi et al., 2020) using Fisher’s Exact test and Bonferroni correction. REVIGO (Supek et al., 2011) was used to reduce GO term redundancy. R environment (R Foundation) and its packages (ggplot2, pheatmap, ComplexHeatmap, dendsort, rstatix, ggpubr, dplyr, reshape2, tidyverse, RColorBrewer, circlize, ggfortify, gridExtra) were used for statistical analysis and to visualize results.

### Ethylene treatment

For ethylene treatments, WT seedlings were grown for 4 days as previously described in unshaded conditions, then transferred to new plates containing 0, 0.2, 2 or 10 μM of ACC. For each treatment, 15 well grown seedlings were transferred for the final plates. Then, they were kept either in control or shaded conditions for 4 days. Plates were then scanned for phenotypic analyses.

### Statistical Analysis

Most statistical analyses were performed in RStudio. Analysis of variance (two-way ANOVA) and pairwise *post-hoc* Tukey analysis were performed in Infostat. Phenotypic data was analyzed by comparing the means between treatments or genotypes according to the test specified at the figures.

### Accession Numbers

WRKY13 (AT4G39410), WRKY25 (AT2G30250), WRKY26 (AT5G07100), WRKY29 (AT4G23550), WRKY31 (AT4G22070), WRKY45 (AT3G01970), WRKY51 (AT5G64810), WRKY58 (AT3G01080), WRKY70 (AT3G56400), WRKY75 (AT5G13080), EIN2 (AT5G03280), EIN3(AT3G20770), EIL1(AT2G27050).

## ACKNOWLEDGEMENTS

We thank Jillian Arber and Jaynee Hart for the technical assistance, cloning, and plant care. We thank Hong Qiao for *ein2* and *ein3eil1* seeds. This work was funded by National Science Foundation (NSF) Division of Integrative Organismal Systems grant IOS-1755355 to U.V.P. National Institutes of Health (NIH) grants R35GM125003, GM12500303S1, and GM12500304S1 to U.V.P. São Paulo Research Foundation (FAPESP) grant no. 2016/01128-9 and the Brazilian National Council of Scientific and Technological Development (CNPq) funded M.R.

## SUPPLEMENTAL FIGURE LEGENDS

**Figure S1. Transcriptional changes induced by shade in the roots.**

A) Phenotype of representative 9-day-old *Arabidopsis* seedlings under constant white light (unshaded) and shade (low R:FR). The phenotype represents 4-day-old seedlings grown in unshaded light and then transferred to the shade or mock-treated for 5 days B) Heatmap of the expression profile of differentially expressed genes (DEG), both up and downregulated that are statistically significant in at least one time-point (FDR <0.05) in *Arabidopsis* and tomato. C) *de novo* enriched cis-motif elements found in the promoters of the 4,835 genes induced, and D) 4,570 repressed by the shade in *Arabidopsis* roots. E) Expression profile of WRKY target genes *CYP71A12, MYB51*, and *PIP1* in response to the shade. Values represent log_2_ FPKM for shade and unshaded control.

**Figure S2. Shade induces the expression of a large group of *WRKYs* in tomato.**

*WRKYs* that are significantly up-regulated (FDR <0.05) in the roots of tomato seedlings grown in the shade relative to its unshaded control during the course of the experiment.

**Figure S3. WRKYox-Citrine protein levels are unaffected by shade.**

Protein expression levels of the indicated WRKYox-Citrine fusions in 4-5 day-old *Arabidopsis* transgenic seedlings. Seedlings were grown in constant white light (unshaded) for 4d and then transferred to the shade for 3h or 24h prior to harvest. WRKY-Citrine proteins were detected with an anti-GFP primary antibody. Ponceau staining of the Rubisco proteins serves as a loading control.

**Figure S4. Phenotype of independent *Arabidopsis* WRKY*ox* transgenic lines in the shade.**

The phenotypic measurement represents 4-day-old seedlings grown in unshaded light and then transferred to the shade or mock-treated for 5 days. Three independent transgenic lines for each indicated WRKY was analyzed. A) Primary root length in cm, B) Hypocotyl length in cm, C) Lateral root number, D) Frequency of seedlings with 0 to 10 or more lateral roots, and E) Lateral root density is the number of lateral roots per cm of the primary root in 9-days old seedlings under unshaded or shade conditions. A-C, E represent means ± SE of individual lines (*UBQ10::WRKYox*) for each candidate gene; dots represent individual data points. Asterisks represent significant differences between transgenics and WT control (black) or within genotypes between shaded and unshaded condition (red) (* *p*<0.5, ** *p*<0.01, *** *p*<0.001) in two-tailed *t*-test with Benjamini-Hochberg correction for multiple testing.

**Figure S5. Overexpression of WRKY13, 29 and 58 does not affect root growth.**

The phenotypic measurement represents 4-day-old seedlings grown in unshaded light and then transferred to the shade or mock-treated for 4 days. Three independent transgenic lines for each indicated WRKY was analyzed. A,D) Primary root length in cm, B,E) frequency of seedlings with 0 to 10 or more lateral roots and C,F) lateral root density in number of lateral roots per cm of primary root under unshaded or shade conditions in 8-days old seedlings. A,C,D and F, values represent means ± SE of transgenic lines (*UBQ10::WRKYox*) for each candidate gene; dots represent individual data points. A-C, combined data from all three transgenic lines; D-F, data of each independent line. Asterisks represent significant differences between transgenics and WT control (black) or within genotypes between shaded and unshaded condition (red) (* *p*<0.5, ** *p*<0.01, *** *p*<0.001) in two-tailed *t*-test with Benjamini-Hochberg correction for multiple testing.

**Supplemental Table S1.**Expression levels (log2 fold enrichment) of genes differentially expressed in the shade relative to its unshaded control (FDR < 0.05) in the roots of *Arabidopsis* and tomato.

**Supplemental Table S2.**Gene Ontology (GO) terms enriched among the shade-induced genes in roots of *Arabidopsis* and tomato.

**Supplemental Table S3.**Oligonucleotide sequences and plasmids used in this work.

## List of author contributions

U.V.P. conceived the study. D.R., M.R. and U.V.P. designed experiments. D.R. performed most of the experiments and analyzed the data; A.A. and O.S. performed genotyping, generation of transgenic plants, and immunoblots, D.R. and U.V.P. wrote the paper and collected contributions of all authors.

U.V.P. agrees to serve as the author responsible for contact in accordance with the journal’s policy.

