## Supplemental Figures for "Shade-induced WRKY transcription factors restrict root growth during the shade avoidance response"

**Supplemental Figure S1, Rosado et al.**

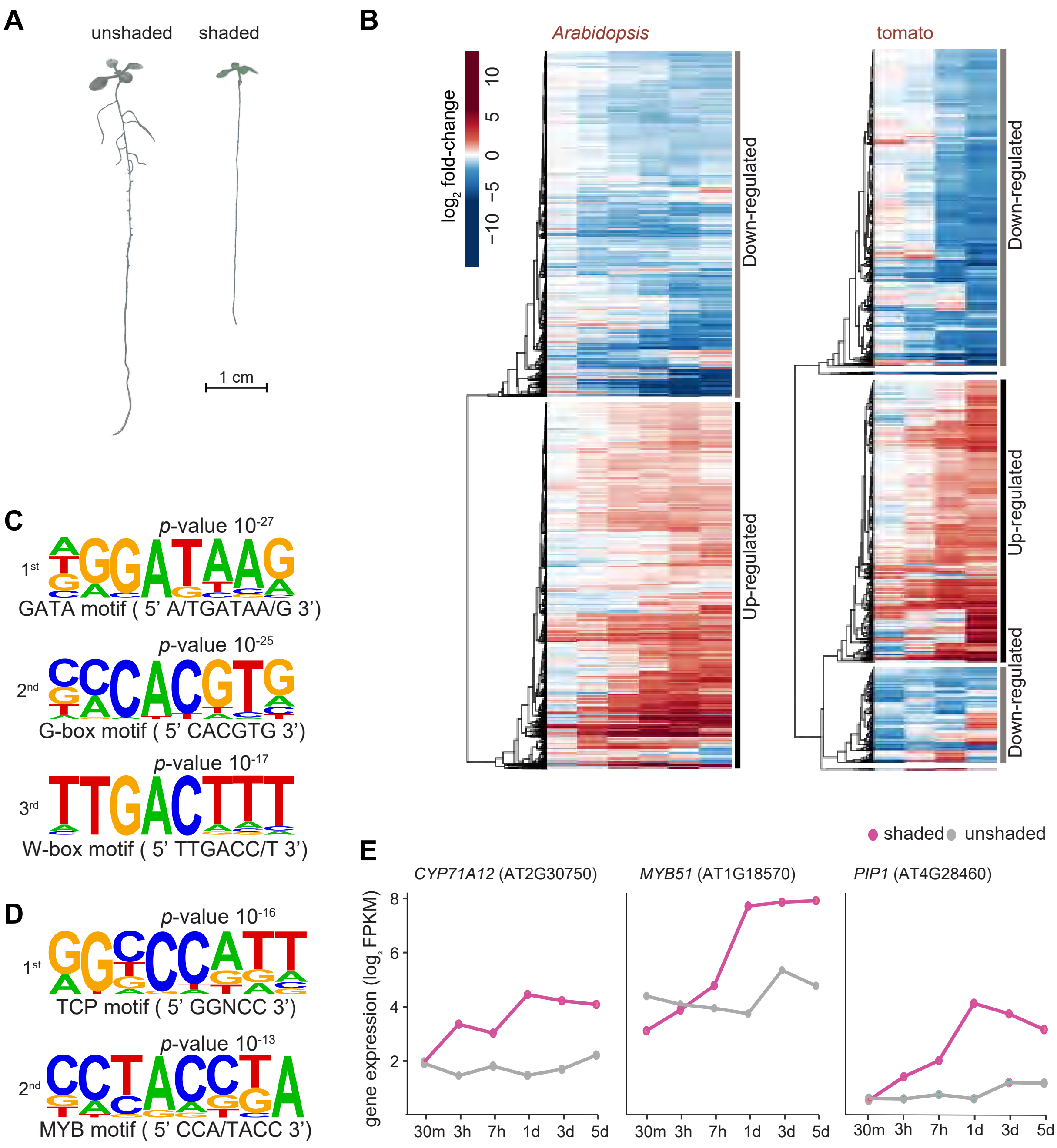


**Fig. S1. Transcriptional changes induced by shade in the roots.**
A) Phenotype of representative 9-day-old *Arabidopsis* seedlings under constant white light (unshaded) and shade (low R:FR). The phenotype represents 4-day-old seedlings grown in unshaded light and then transferred to the shade or mock-treated for 5 days. B) Heatmap of the expression profile of differentially expressed genes (DEG), both up and downregulated that are statistically significant in at least one time-point (*FDR* <0.05) in *Arabidopsis* and tomato. C) *de novo* enriched *cis*-motif elements found in the promoters of the 4,835 genes induced, and D) 4,570 repressed by the shade in *Arabidopsis* roots. E) Expression profile of WRKY target genes *CYP71A12*, *MYB51*, and *PIP1* in response to the shade. Values represent $\log_2$ FPKM for shade and unshaded control.

**Supplemental Figure S2, Rosado et al.**

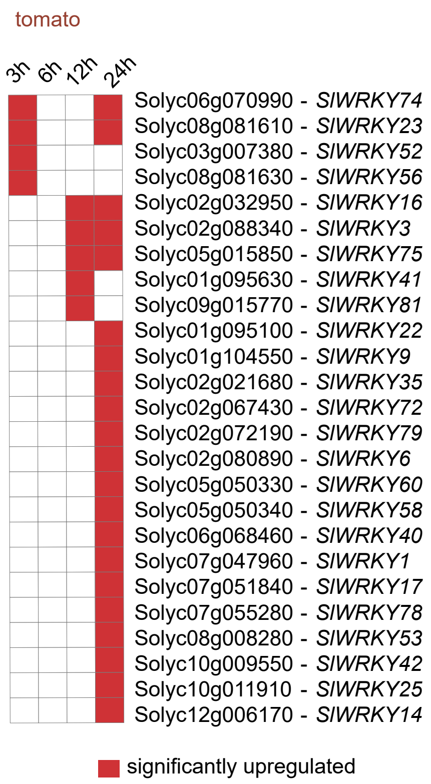


**Fig. S2. Shade induces the expression of a large group of WRKYs in tomato.**
*WRKYs* that are significantly up-regulated (FDR <0.05) in the roots of tomato seedlings grown in the shade relative to its unshaded control during the course of the experiment.

**Supplemental Figure S3, Rosado et al.**

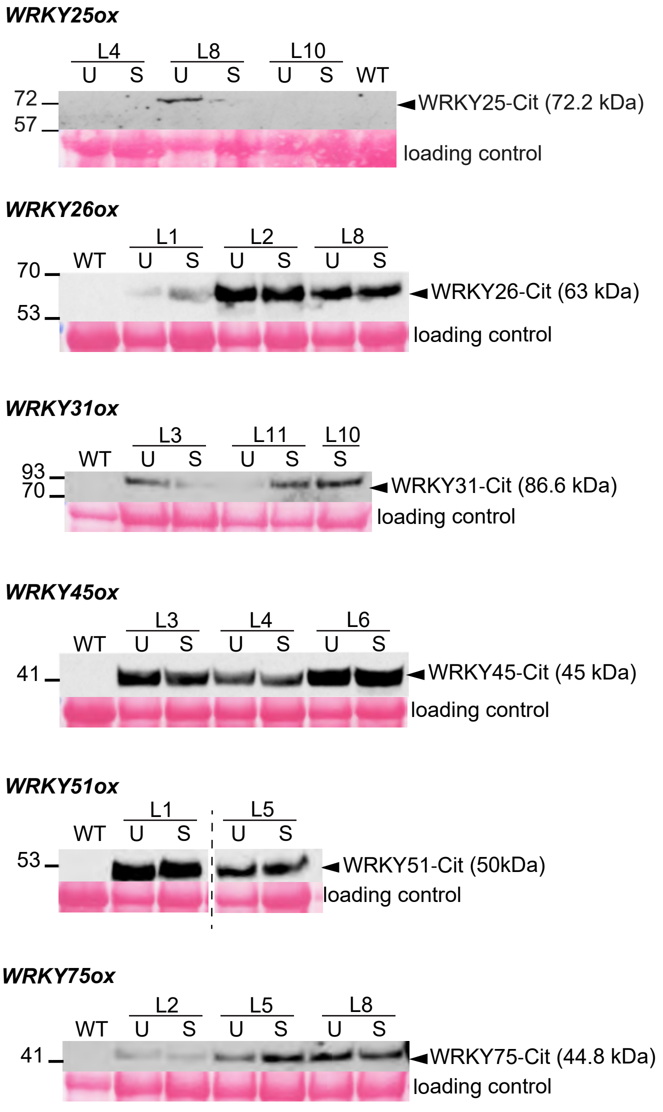


**Fig. S3. WRKYox-Citrine protein levels are unaffected by shade.**
Protein expression levels of the indicated WRKYox-Citrine fusions in 4-5 day-old *Arabidopsis* transgenic-seedlings. Seedlings were grown in constant white light (unshaded) for 4d and then transferred to the shade for 3h or 24h prior to harvest. WRKY-Citrine protein levels were detected with an anti-GFP primary antibody. Ponceau staining of the Rubisco proteins serves as a loading control.

### Supplemental Figure S4, Rosado et al.

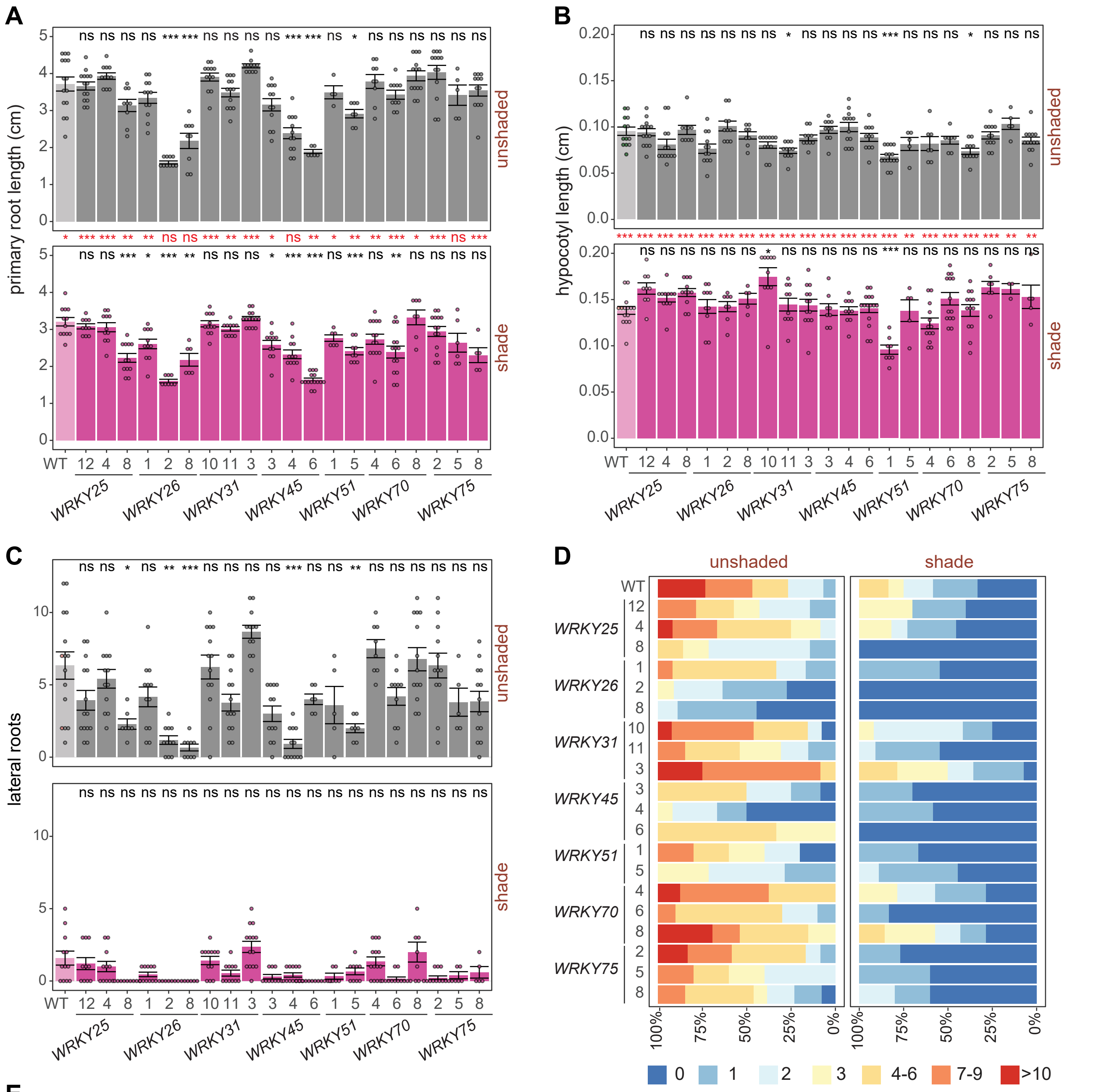


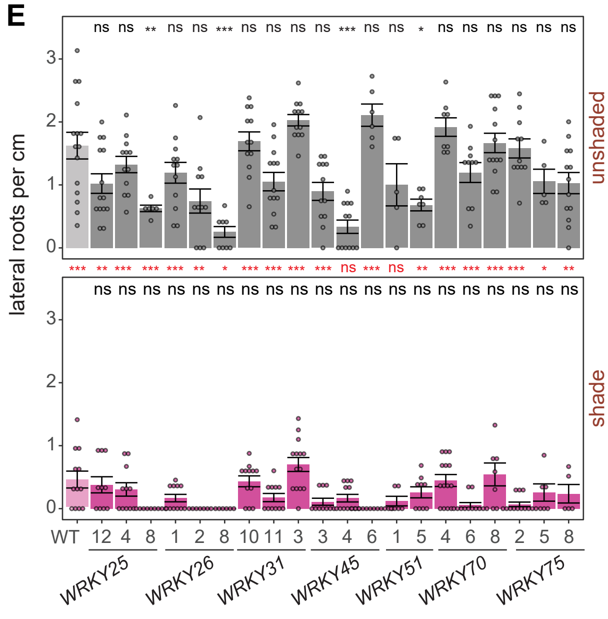


**Fig. S4. Phenotype of independent *Arabidopsis* WRKY*ox* transgenic lines in the shade.**
The phenotypic measurement represents 4-day-old seedlings grown in unshaded light and then transferred to the shade or mock-treated for 5 days. Three independent transgenic lines for each indicated WRKY was analyzed. A) Primary root length in cm, B) Hypocotyl length in cm, C) Lateral root number, D) Frequency of seedlings with 0 to 10 or more lateral roots, and E) Lateral root density is the number of lateral roots per cm of the primary root in 9-days old seedlings under unshaded or shade conditions. A-C, E represent means ± SE of individual lines (*UBQ10::WRKYox*) for each candidate gene; dots represent individual data points. Asterisks represent significant differences between transgenics and WT control (black) or within genotypes between shaded and unshaded condition (red) (* $p$<0.5, ** $p$<0.01, *** $p$<0.001) in two-tailed $t$-test with Benjamini-Hochberg correction for multiple testing.

### Supplemental Figure S5, Rosado et al.

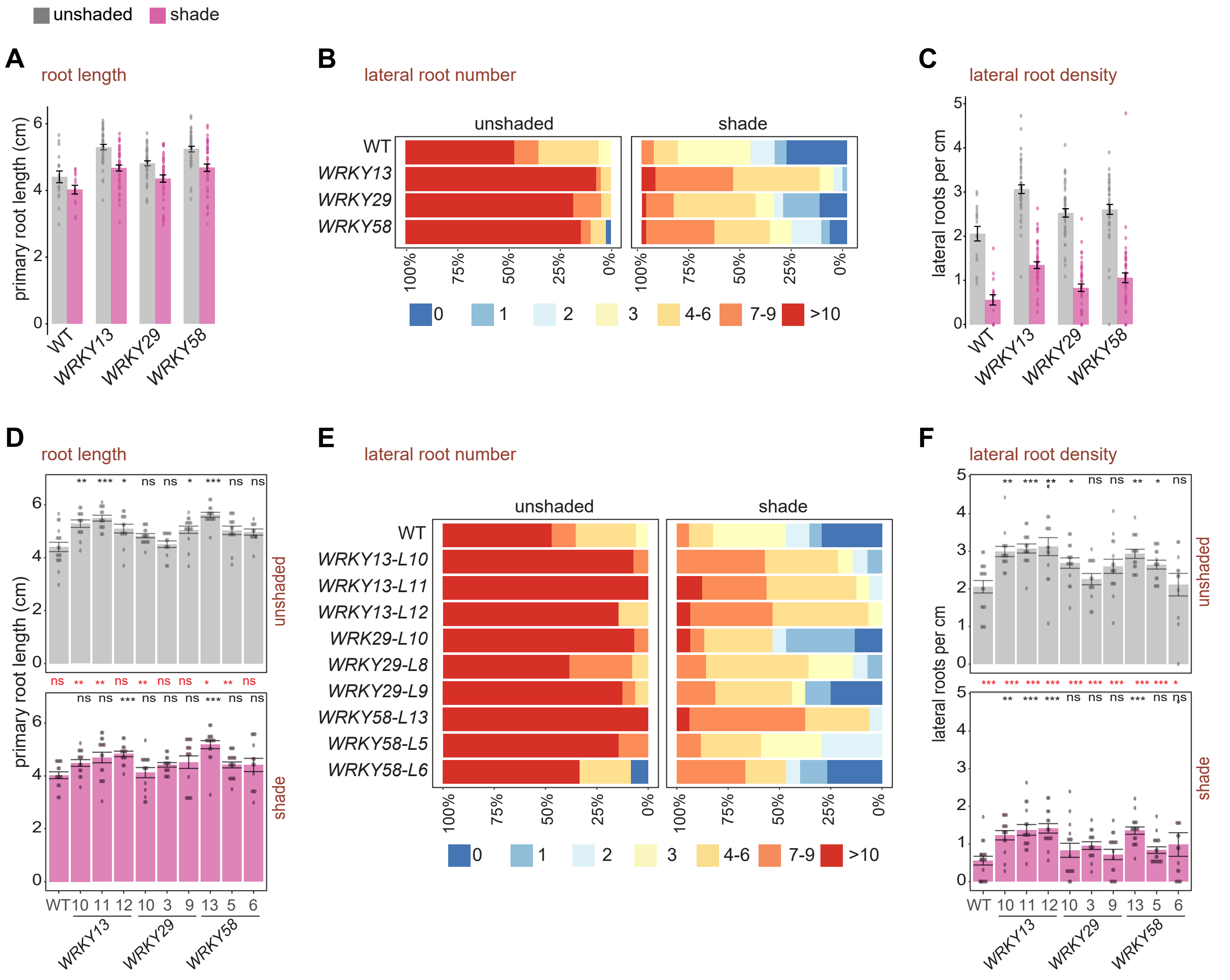


**Figure S5. Overexpression of WRKY13, 29 and 58 does not affect root growth.**
The phenotypic measurement represents 4-day-old seedlings grown in unshaded light and then transferred to the shade or mock-treated for 4 days. Three independent transgenic lines for each indicated WRKY was analyzed. A,D) Primary root length in cm, B,E) frequency of seedlings with 0 to 10 or more lateral roots and C,F) lateral root density in number of lateral roots per cm of primary root under unshaded or shade conditions in 8-days old seedlings. A,C,D and F, values represent means ± SE of transgenic lines (*UBQ10::WRKYox*) for each candidate gene; dots represent individual data points. A-C, combined data from all three transgenic lines; D-F, data of each independent line. Asterisks represent significant differences between transgenics and WT control (black) or within genotypes between shaded and unshaded condition (red) (* *p*<0.5, ** *p*<0.01, *** *p*<0.001) in two-tailed *t*-test with Benjamini-Hochberg correction for multiple testing.
